# Functional maturation of the gut microbiota at weaning is influenced by maternal environment in piglets

**DOI:** 10.1101/2020.04.02.022913

**Authors:** Martin Beaumont, Laurent Cauquil, Allan Bertide, Ingrid Ahn, Céline Barilly, Lisa Gil, Cécile Canlet, Olivier Zemb, Géraldine Pascal, Arnaud Samson, Sylvie Combes

## Abstract

The objective of this study was to analyze in piglets the impact of weaning on the production of metabolites by gut bacteria and to determine whether early life environment influences the functional maturation of the gut microbiota. Fecal metabolome and microbiome were analyzed in piglets raised in two separate maternity farms and mixed at weaning. In piglets from both maternity farms, the relative abundance of *Lactobacillus* and of the predicted function “Fucose degradation” decreased after weaning while the relative abundance of *Ruminococcus 2* and of the predicted function “Starch degradation” increased. In piglets from the first maternity farm, the relative concentration of biogenic amines and the relative abundance of *Escherichi-Shigella* decreased after weaning while the relative concentration of short chain fatty acids and the relative abundance of *Christensenellaceae R-7 group* and *Ruminococcaceae UCG-002* increased. These changes were not observed at weaning in piglets from the second maternity farm probably because they already had high relative concentration of short chain fatty acids and higher relative abundance of *Christensenellaceae R-7 group* and *Ruminococcaceae UCG-002* during the suckling period. In conclusion, the functional maturation of the microbiota at weaning is highly dependent on the maternal environment in piglets.

**ORIGINALITY - SIGNIFICANCE STATEMENT:** Bacterial metabolites are key molecular intermediates between the gut microbiota and host cells. Our study in piglets reveals that the metabolic activity of the gut microbiota shifts at weaning, a key developmental period for intestinal and immune health. We also show that this functional maturation of the gut microbiota is strongly influenced by maternal environment. Thus, targeting early life environmental factors is a promising strategy to program health trough the production of beneficial bacterial metabolites at the suckling-to-weaning transition.

## INTRODUCTION

The gut microbiota is a major regulator of animal physiology and health. After initial colonization at birth by microorganisms originating both from the mother and the environment, maternal milk shapes the gut microbiota composition through nutrients (e.g. lactose, milk oligosaccharides), immunoglobulins and antimicrobial compounds (e.g. lactoferrin, lysozyme) (Macpherson *et al.*, 2017). Later in life, solid food ingestion induces the maturation of the gut microbiota mainly through a modification of dietary substrates available for bacteria (Voreades *et al.*, 2014). These microbial modifications at the suckling-to-weaning transition are involved in the postnatal maturation of the gut barrier and immune system (Hooper, 2004; Jain and Walker, 2015). Moreover, it was recently shown that these alterations of the gut microbiota at the onset of solid food ingestion induce a transient intestinal immune response called the “weaning-reaction” that programs long term susceptibility of the host to inflammatory and metabolic dysfunctions (Al Nabhani, Dulauroy, Lécuyer, *et al.*, 2019; Al Nabhani, Dulauroy, Marques, *et al.*, 2019). Although the underlying mechanisms are not fully identified yet, emerging evidences suggest that bacterial metabolites may play a key role as intermediates between the microbiota and its host at the suckling-to-weaning transition (Al Nabhani and Eberl, 2020). Therefore, identifying the metabolites produced by gut bacteria in young mammals across this dietary shift might be useful for the development of health-promoting strategies based on the control of the metabolic activity of the microbiota in early life.

Piglets raised in commercial conditions represent an attractive animal model to study the effects of weaning on the microbiota since the separation from the sow occurs 3 to 4 weeks after birth (versus 8 to 14 weeks in natural conditions) which results in an abrupt cessation of suckling when piglets milk intake is still high while solid feed intake very low (Newberry and Wood-Gush, 1985). Moreover, the results obtained in this model are relevant both for the pig industry (e.g. management of post-weaning diarrhea in piglets) and for human health due to the anatomical, functional and microbial similarities between piglet and human infant gastrointestinal tract (Heinritz *et al.*, 2013; Gresse *et al.*, 2017). Numerous studies in piglets described a major shift in the microbiota taxonomic composition and functional capacity at the suckling-to-weaning transition (Frese *et al.*, 2015; Mach *et al.*, 2015; Slifierz *et al.*, 2015; Bian *et al.*, 2016; Chen *et al.*, 2017; De Rodas *et al.*, 2018; Guevarra *et al.*, 2018; Li *et al.*, 2018; Lu *et al.*, 2018; W. Wang *et al.*, 2019; X. Wang *et al.*, 2019). Although to a much lower extent, maternal environment (e.g. nursing mother) was also shown to impact the microbiota composition in piglets around weaning (Bian *et al.*, 2016). However, these studies were limited to DNA sequencing-based approaches and did not investigate the influence of early life environment and weaning on the actual production of metabolites by gut bacteria.

Herein, we analyzed the metabolic activity of the gut microbiota during the suckling period and after weaning in piglets raised in two distinct maternal environments and mixed at weaning in the same room. By using a combination of H^1^-nuclear magnetic resonance (NMR) based metabolomics, 16S rRNA gene sequencing and bacterial pathway inference, our study reveals that the functional maturation of the gut microbiota at weaning is strongly influenced by the maternal environment in piglets.

## RESULTS

### Maternal environment and weaning influence piglet fecal metabolome

We studied piglets raised in two separate maternity farms and mixed at weaning (day 21) in the same pens in one room (figure 1). Metabolome was analyzed by H^1^-NMR metabolomics in fecal samples collected from piglets during the suckling period (day 13) and two days after weaning (day 23). We identified thirty-nine metabolites in the NMR spectra of piglet fecal samples (table 1 and figure 2). During the suckling period (day 13), partial least-square discriminant analysis (PLS-DA) suggested differences of metabolome between piglets from the two maternity farms (figure 3A). Indeed, succinate and 3-(4-hydroxyphenyl)propionate relative concentrations were significantly higher in piglets from maternity 1 compared to piglet from maternity 2, while the opposite was observed for isobutyrate and propionate (figure 3B, table S2). In piglets from maternity 1, we observed a strong modification of fecal metabolome after weaning (figure 3A). Indeed, the relative concentration of cadaverine, tyramine, succinate, 3-(4-hydroxyphenyl)propionate, 5-aminovalerate and choline decreased after weaning in piglets from maternity 1, while the relative concentrations of acetate, propionate, dihydroxyacetone, glycerol, glutamate and uracil increased (figure 3B, table S2). In contrast, in piglets from maternity 2, the metabolic shift observed at weaning was less pronounced according to the PLS-DA analysis (figure 3A). Accordingly, we found no significant effect of weaning on the relative concentrations of individual metabolites in piglets from maternity 2 (figure 3B, table S2). At day 23, the relative concentration of only one metabolite (glycerol) was significantly different between the piglets from the two maternity farms (figure 3B, table S2). Altogether, our results show that the metabolic alteration observed in feces at weaning is influenced by the maternal environment in piglets. Since most of the metabolites which concentration was altered at weaning are known to be produced by bacteria (e.g. biogenic amines, short chain fatty acids (SCFA), succinate, 3-(4-hydroxyphenyl)propionate), our data indicated that the metabolic activity of the gut microbiota shifted at weaning in an early-life environment dependent manner.

**Table 1:** Metabolites identified by NMR metabolomics in piglet fecal samples. *: indicates the peak used for quantification based on the corresponding bucket intensity (not overlapping with peaks from other metabolites). Multiplicity of signals is indicated within brackets: s, singlet; d, doublet; dd, doublet of doublet; t, triplet; q, quadruplet and m, multiplet.

| | Metabolite | $\delta^1\text{H}$ (ppm) |
| --- | --- | --- |
| 1 | 2-methylbutyrate | 0.86* (t), 1.05 (d), 1.39 (m), 1.49 (m), 2.1 (m) |
| 2 | Valerate | 0.89 (t), 1.30* (m), 1.53 (m), 2.19 (t) |
| 3 | Butyrate | 0.90* (t), 1.56 (m), 2.16 (t) |
| 4 | Isovalerate | 0.92 (d), 1.96 (m), 2.06* (d) |
| 5 | 4-methyl-2-oxovalerate | 0.94 (d), 2.10 (m), 2.62* (d) |
| 6 | Isoleucine | 0.94 (t), 1.01* (d), 1.27 (m), 1.47 (m), 1.99 (m), 3.68 (d) |
| 7 | Leucine | 0.97* (t), 1.72 (m), 3.74 (m) |
| 8 | Valine | 1.00* (d), 1.05 (d), 2.28 (m), 3.62 (d) |
| 9 | Propionate | 1.06* (t), 2.19 (m) |
| 10 | Isobutyrate | 1.07 (d), 2.39* (m) |
| 11 | 3-methyl-2-oxovalerate | 0.90 (t), 1.10* (d), 1.46 (m), 1.71 (m), 2.94 (m) |
| 12 | 3-methyl-2-oxobutyrate | 0.90 (d), 1.13* (d), 3.03 (m) |
| 13 | Threonine | 1.34* (d), 4.26 (m) |
| 14 | Cadaverine | 1.48 (m), 1.73* (m), 3.02 (t) |
| 15 | 5-aminovalerate | 1.65 (m), 2.24* (t), 3.02 (t) |
| 16 | Putrescine | 1.78* (m), 3.06 (m) |
| 17 | Acetate | 1.92* (s) |
| 18 | Glutamate | 2.07 (m), 2.14 (m), 2.35* (m), 3.77 (m) |
| 19 | Succinate | 2.41* (s) |
| 20 | Methylamine | 2.60* (s) |
| 21 | Methionine | 2.14 (s), 2.14 (m), 2.20 (m), 2.65* (t), 3.87 (m) |
| 22 | Dimethylamine | 2.72* (s) |
| 23 | Trimethylamine | 2.88* (s) |
| 24 | Tyramine | 2.94 (t), 3.24 (t), 6.92 (d), 7.23* (d) |
| 25 | Choline | 3.21* (s), 3.53 (m), 4.07 (m) |
| 26 | Glucose | 3.26 (m), 3.39 – 3.56 (m), 3.70 – 3.91 (m), 5.24* (d) |
| 27 | Phenylacetate | 3.54* (s), 7.31 (m), 7.38 (m) |
| 28 | Glycine | 3.57* (s) |
| 29 | Glycerol | 3.56 (dd), 3.66* (dd), 3.79 (m) |
| 30 | Ribose | 3.53 (m), 3.62 (dd), 3.66 (m), 3.69 (dd), 3.76 (dd), 3.80-4.02 (m),<br>4.11* (t), 4.14 (m), 4.22 (m), 5.26 (s), 5.39 (d) |
| 31 | Dihydroxyacetone | 4.42* (s) |
| 32 | Galactose | 3.50 (dd), 3.66 (dd), 3.69-3.88 (m), 3.94 (d), 4.00 (d), 4.09 (m),<br>5.27* (d) |
| 33 | Uracil | 5.81* (d), 7.56 (d) |
| 34 | 3-(4-hydroxyphenyl)propionate | 2.45 (t), 2.81 (t), 6.86* (d), 7.18 (d) |
| 35 | Tyrosine | 3.07 (dd), 3.20 (dd), 3.95 (dd), 6.90 (d), 7.20* (d) |
| 36 | Phenylalanine | 3.14 (dd), 3.29 (dd), 4.01 (dd), 7.33 (d), 7.38 (t), 7.43* (t) |
| 37 | Benzoate | 7.49* (t), 7.56 (t), 7.88 (d) |
| 38 | Hypoxanthine | 8.20* (s), 8.22 (s) |
| 39 | Formate | 8.46* (s) |

**Figure 1:**
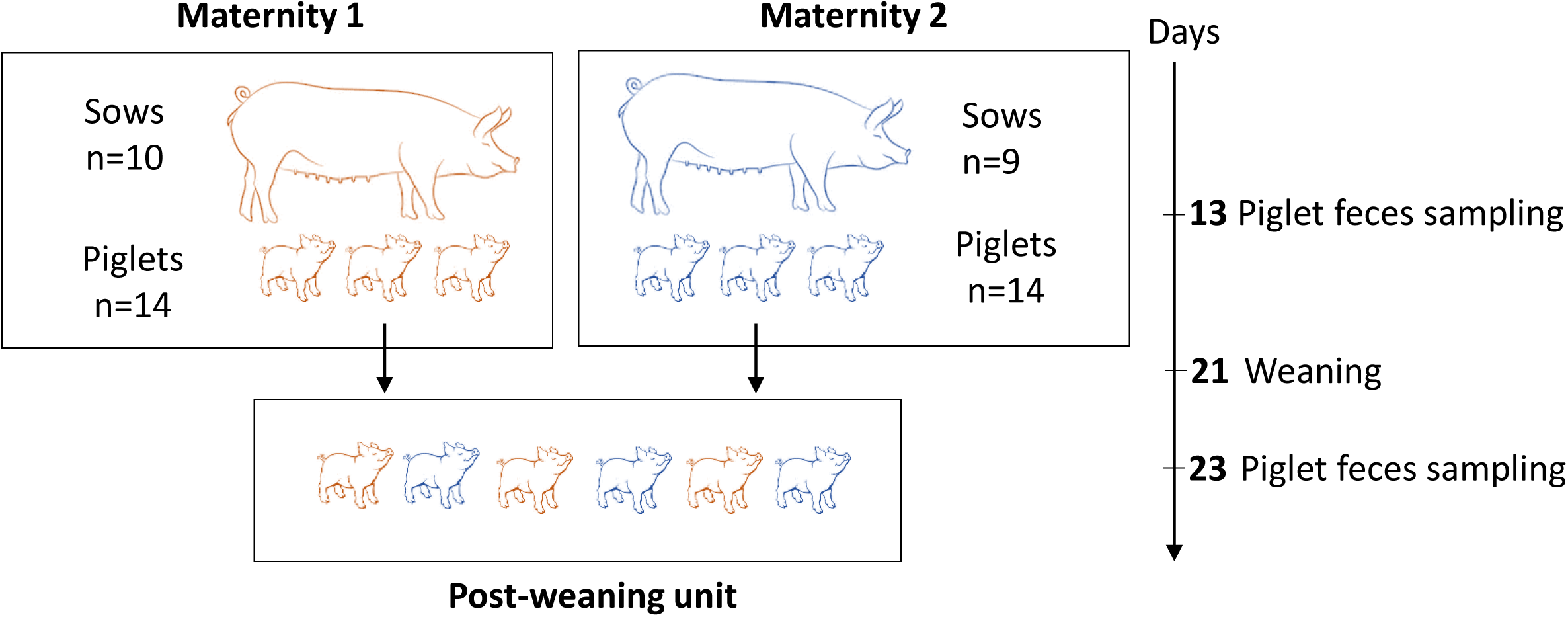
Schematic representation of the experimental design. Piglets from two separate maternity farms (n=14 piglets/maternity) were mixed at weaning (21 days after birth) in the same pens in one room. Fecal samples were collected from the same piglets during the suckling period (day 13) or 2 days after weaning (day 23).

**Figure 2:**
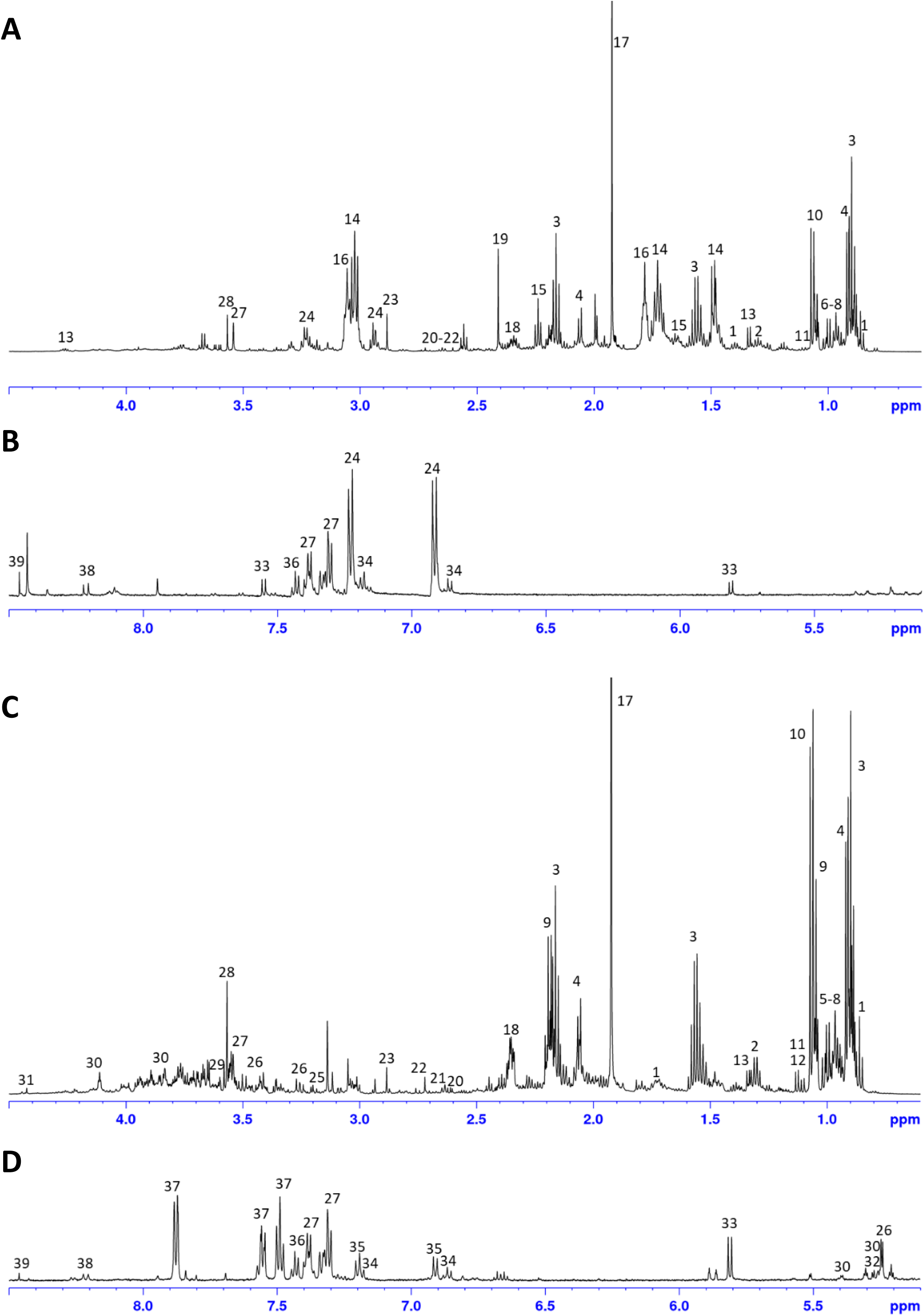
Identification of metabolites in piglets feces NMR spectra. Piglet fecal metabolome was analyzed by NMR metabolomics. A & B: Representative spectra during the suckling period (day 13). C & D: Representative spectra after weaning (day 23). The aliphatic (A, C) and aromatic (B, D – vertically expanded) regions are shown. Peaks are identified with a number corresponding to the metabolites described in table 1.

**Figure 3:**
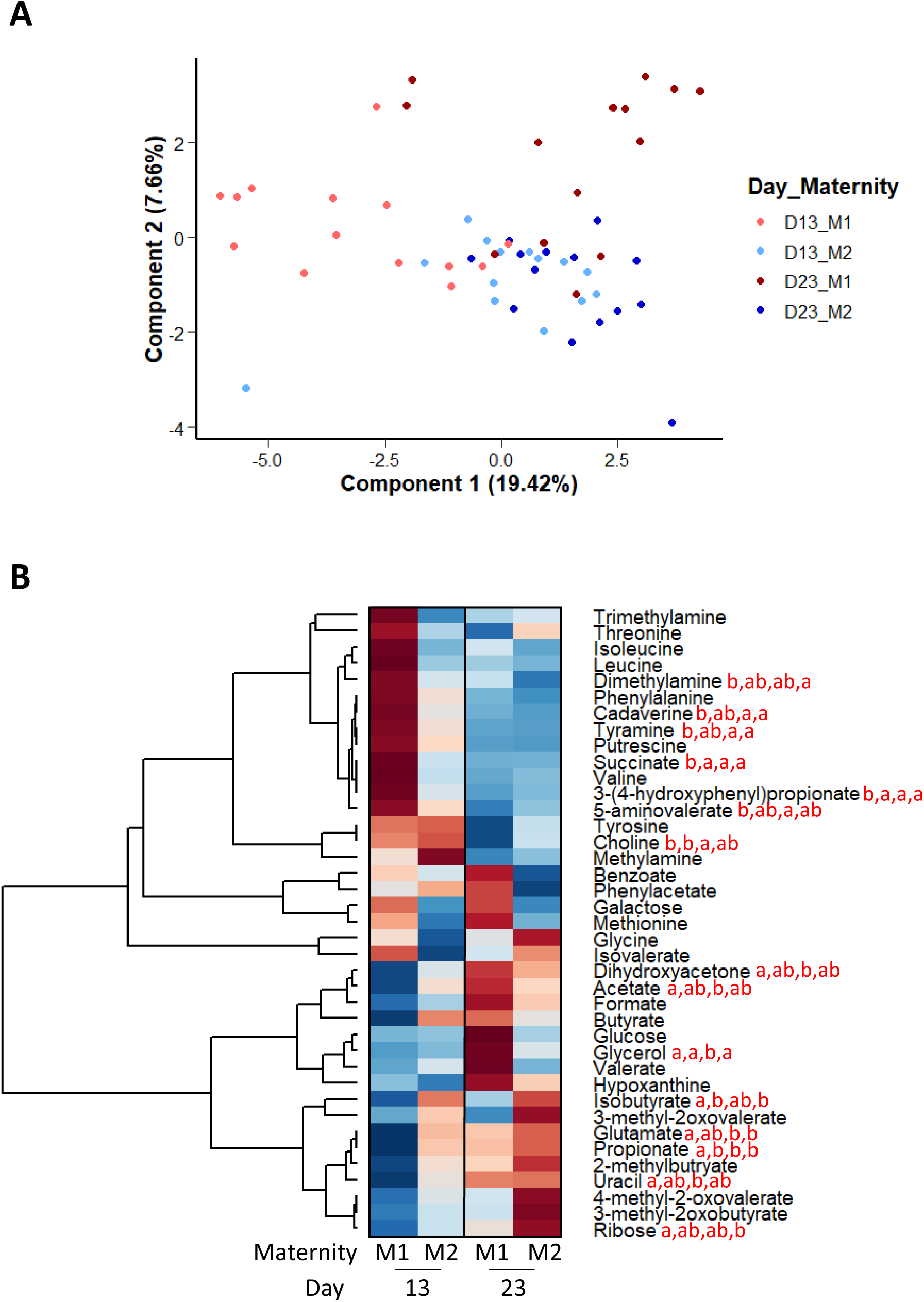
Maternal environment and weaning influence the metabolic activity of the gut microbiota. Metabolome was analyzed by NMR-based metabolomics in feces collected from piglets from two maternity farms (M1 and M2) during the suckling period (D13) and after weaning (D23). A – Individual plot of partial least square discriminant analysis (PLS-DA) with the relative concentration of metabolites used as variable matrix and groups as predictors. B – Heatmap representing the mean relative concentration of each metabolites in each group. The color represent the Z-scores (row-scaled relative concentration) from low (blue) to high values (red). Metabolites (rows) were clustered by the average method. A linear mixed model was used with age and maternity as fixed effects and sows and piglets as random effects. The means of groups associated with different letters are significantly different.

### Maternal environment and weaning influence piglet fecal microbiota diversity and composition

We explored whether the functional maturation of the gut microbiota at weaning was linked to a modification of its taxonomic composition by using 16S rRNA amplicon sequencing in the same fecal samples than those used for metabolomics analysis. At day 13, there was no significant difference between the α-diversity of the microbiota of piglets from the two maternity farms (figure 4A, table S3). In piglets from maternity 1, the three diversity indices tested (OTU richness, Shannon and Inverse Simpson) increased significantly after weaning. In contrast, in piglets from maternity 2, there was only a significant increase of OTU richness after weaning, while the two other diversity indices remained unchanged. At day 23 (i.e. after weaning), there was no significant differences in α-diversity between the piglets from the two maternity farms. β-diversity analysis at OTU level using the Bray-Curtis distance revealed a strong effect of both maternal environment and of weaning on the microbiota structure (figure 4B).

**Figure 4:**
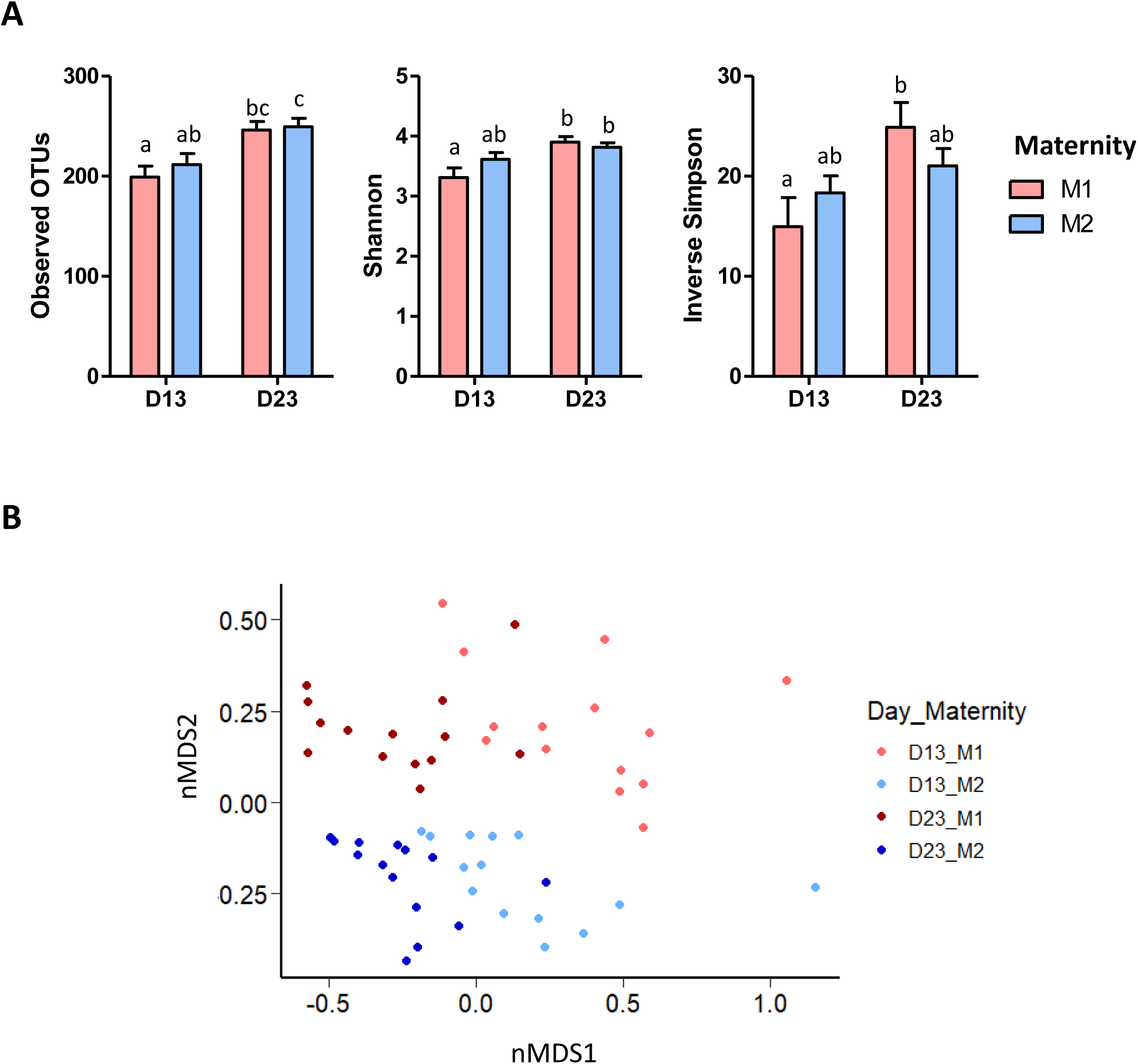
Maternal environment and weaning influence diversity and structure of the gut microbiota. The microbiota composition was analyzed by 16S rRNA gene sequencing in feces collected from piglets from two maternity farms (M1 and M2) during the suckling period (D13) and after weaning (D23). A - α-diversity indices (Mean + SEM). A linear mixed model was used with age and maternity as fixed effects and sows and piglets as random effects. The means of groups associated with different letters are significantly different. B - Non Metric Dimensional Scaling (nMDS) two-dimensional representation of the microbiota β-diversity using Bray Curtis distance calculation (stress=17.68).

The relative abundances of bacterial groups are presented at phylum (table S4), family (figure 5A and table S5), genus (figure 5B for the top 10 most abundant groups and table S6) and OTU level (table S7). At day 13, the relative abundances of Clostridiales vadinBB60 group, Christensenellaceae, *Christensenellaceae R-7 group* and *Ruminococcaceae UCG-002* were higher in piglets from maternity 2 compared to piglets from maternity 1. After weaning, in piglets from both maternities, there was an increase in the relative abundance of Ruminococcaceae and *Ruminococcus 2* and a decrease in the relative abundance of Lactobacillaceae and *Lactobacillus*. In addition, some effects of weaning were observed only in piglets from one of the two maternity farms studied. In piglets from maternity 1, there was a reduction of the relative abundance of Proteobacteria, Enterobacteriaceae and *Escherichia-Shigella* after weaning while the relative abundances of Christensenellaceae, *Christensenellaceae R-7 group* and *Ruminococcaceae UCG-002* increased. In piglets from maternity 2, the relative abundance of Peptostreptococcaceae increased after weaning. After weaning (at day 23), the relative abundances of Spirochaetes and *Ruminococcaceae UCG-002* were higher in piglets from maternity 2 when compared to piglets from maternity 1 while the opposite was observed for Atopobiaceae. Altogether, these results show that both maternal environment and weaning shape the microbiota composition in piglets.

**Figure 5:**
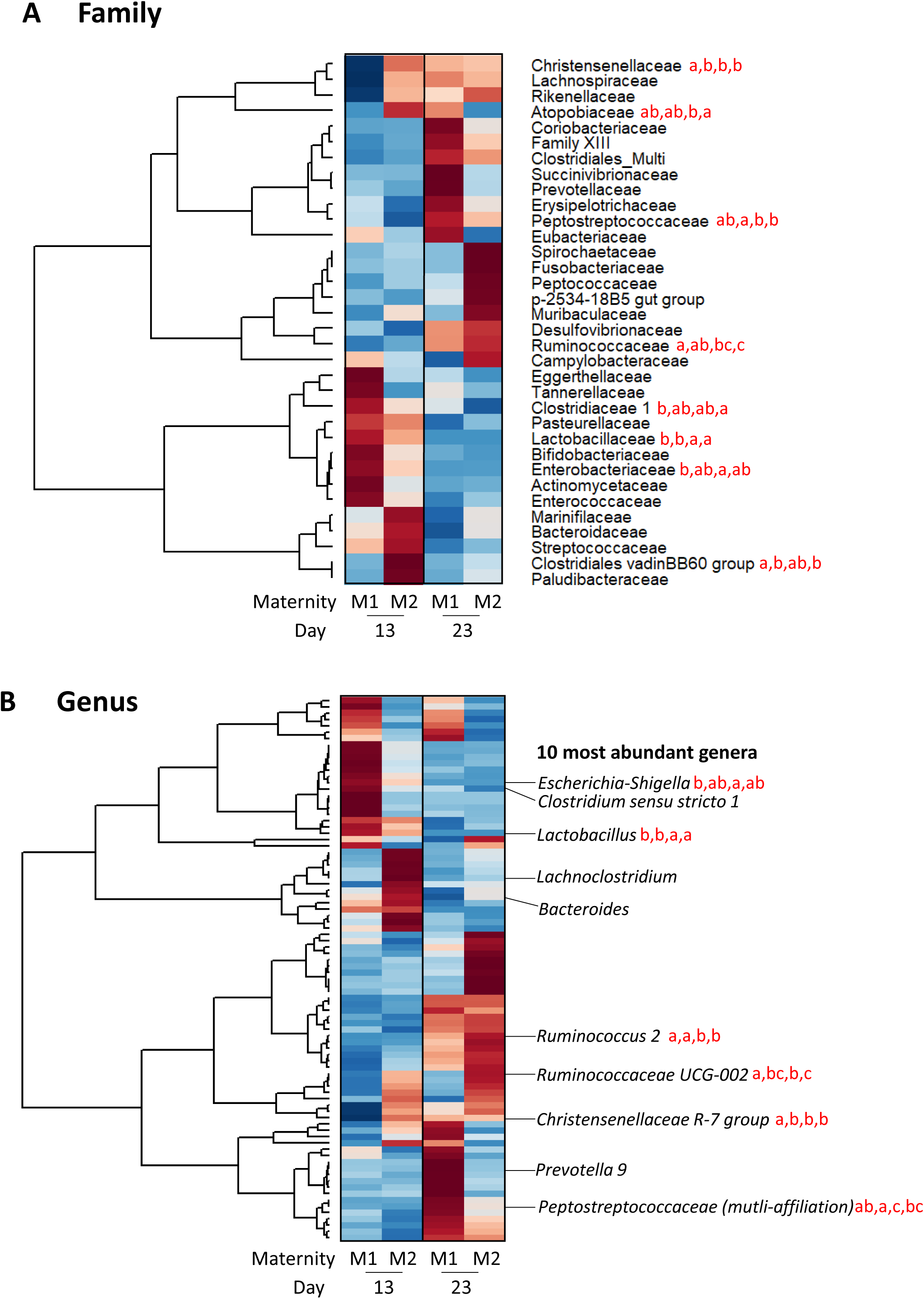
Maternal environment and weaning influence the composition of the gut microbiota. The microbiota composition was analyzed by 16S rRNA gene sequencing in feces collected from piglets from two maternity farms (M1 and M2) during the suckling period (D13) and after weaning (D23). A - Heatmap representing the mean relative abundance of bacterial families in each group. The color represent the Z-scores (row-scaled relative abundance) from low (blue) to high values (red). Families (rows) were clustered by the average method. B - Heatmap representing the mean relative abundance of bacterial genera in each group. The color represent the Z-scores (row-scaled relative abundance) from low (blue) to high values (red). Genus (rows) were clustered by the average method. The names of the 10 most abundant genera are indicated. After log transformation of bacterial groups relative abundances, a linear mixed model was used with age and maternity as fixed effects and sows and piglets as random effects. The means of groups associated with different letters are significantly different.

### Maternal environment and weaning influence the predicted functionality of the gut microbiota

We predicted the functional capacity of gut bacteria by using inference based on 16S rDNA amplicons sequences (PICRUSt2). Heatmap representation of the relative abundance of all predicted pathways suggested that the functional potential of the microbiota was influenced mainly by age and to a lower extent by maternal environment (figure 6A). Indeed, there was a significant effect of weaning for 62% of the predicted pathways (199/320) and a significant effect of maternal environment for only 5% of them (15/320) (table S8). We focused on two key microbial pathways involved in the degradation of dietary substrates available for the gut microbiota: fucose (a monosaccharide contained in milk oligosaccharides) and starch (a major plant carbohydrate).

**Figure 6:**
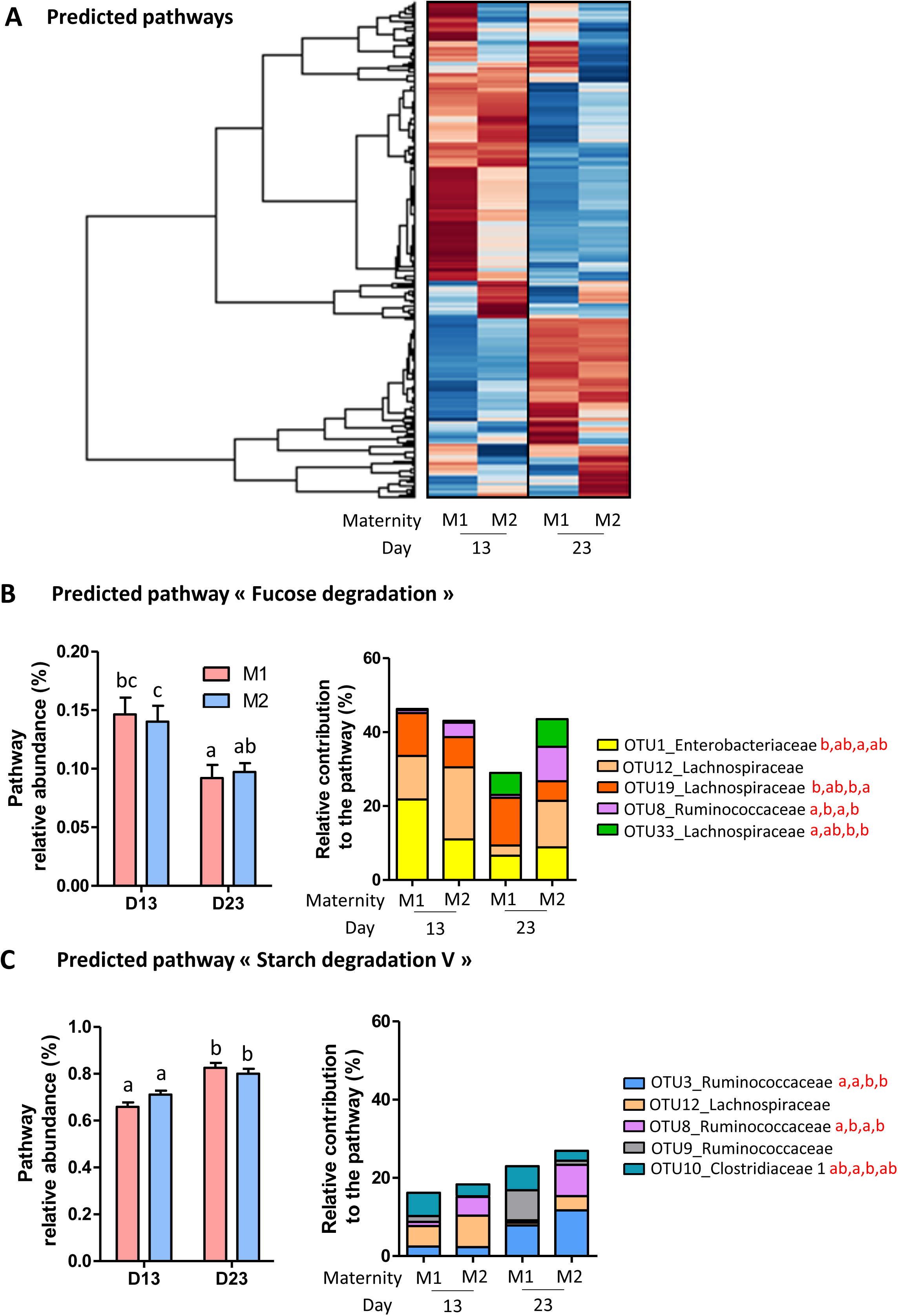
Maternal environment and weaning influence the predicted functionality of the gut microbiota. The potential functionality of the gut microbiota was inferred from 16S rRNA amplicon sequences in feces collected from piglets from two maternity farms (M1 and M2) during the suckling period (D13) and after weaning (D23). A – Heatmap representing the mean relative abundance of each predicted pathways in each group. The color represent the Z-scores (row-scaled relative abundance) from low (blue) to high values (red). Predicted pathways (rows) were clustered by the average method. B – “Fucose degradation” predicted pathway. C – “Starch degradation V” predicted pathway. Left panels: relative abundance of the predicted pathway (mean + SEM). Right panels: relative contribution of the 5 OTUs contributing the most to the predicted pathway (means). After log transformation of bacterial predicted pathway relative abundances or OTU relative contribution to the pathway, a linear mixed model was used with age and maternity as fixed effects and sows and piglets as random effects. The means of groups associated with different letters are significantly different.

The relative abundance of the predicted pathway “Fucose degradation” decreased after weaning in piglets from both maternity farms (figure 6B). The 5 OTUs contributing the most to this predicted function were affiliated to Enterobacteriaceae, Lachnospiraceae and Ruminococcaceae (figure 6B and table S9). At day 13, the relative contribution of OTU8 (Ruminococcaceae) to this function was more important in piglets from maternity 2 than in piglets from maternity 1. In piglets from maternity 1, the relative contribution of OTU1 (Enterobacteriaceae) to the fucose degradation pathway decreased after weaning while the opposite was observed for the OTU33 (Lachnospiraceae). In piglets from maternity 2, the relative contribution of the 5 OTUs contributing the most to this pathway did not change after weaning. At day 23, the relative contribution of OTU8 (Ruminococcaceae) was still more important in piglets from maternity 2 compared to piglets from maternity 1 while the opposite was observed for the relative contribution of OTU19 (Lachnospiraceae).

The relative abundance of the predicted pathway “Starch degradation V” increased after weaning in piglets from both maternity farms (figure 6C). The 5 OTUs contributing the most to this function were affiliated to Ruminococcaceae, Lachnospiraceae and Clostriadiaceae 1 (figure 6C and table S10). At day 13, the relative contribution of OTU8 (Ruminococcaceae) to this function was more important in piglets from maternity 2 compared to piglets from maternity 1. The relative contribution of OTU3 (Ruminococcaceae) increased after weaning in piglets from both maternity farms. At day 23, the relative contribution of OTU8 (Ruminococcaceae) to this function was still more important in piglets from maternity 2 compared to piglets from maternity 1 (as observed at day 13).

It is important to consider that the relative contribution of OTUs to these two predicted pathways was strongly and positively correlated with the OTUs relative abundances (table S7, ρ>0.86 for both pathways). Moreover, all the OTUs contributing the most to the predicted pathways had a high relative abundance (>1.4% in at least one group, table S7). Overall, our results show that weaning and, to a lower extent, maternal environment influence the predicted functionality of the gut microbiota and the relative contributions of bacterial OTUs to these functions.

## DISCUSSION

Our study reveals that the production of metabolites by gut bacteria shifts at weaning in piglets in an early life environment-dependent manner. Since bacterial metabolites which concentration was altered at weaning (e.g. SCFA and biogenic amines) are known to regulate the homeostasis of host cells (Koh *et al.*, 2016; Tofalo *et al.*, 2019), the metabolic shift of the microbiota observed at weaning might have major implication for health. Moreover, the influence of maternal environment on the functional maturation of the microbiota at weaning in piglets suggests that controlling early life environment might be a promising strategy to promote the production of beneficial metabolites at weaning, and thus to program long term health (Al Nabhani and Eberl, 2020).

During the suckling period, the microbiota of piglets raised in the two separate maternity farms produced different levels of several bacterial metabolites. For instance, the relative concentration of 3-(4-hydroxyphenyl)propionate was higher in piglets from maternity 1 during the suckling period. This metabolite is produced by the microbiota through degradation of the aromatic amino acid tyrosine (Oliphant and Allen-Vercoe, 2019) which concentration was similar in the feces of piglets from both maternities. Thus, rather than substrate availability, differences in microbiota composition and functional potential might drive the differential production of 3-(4-hydroxyphenyl)propionate. We also observed a high concentration of succinate in fecal samples of piglets from maternity 1, despite this metabolic intermediate usually do not accumulate since it can be rapidly converted to SCFA by gut bacteria (Oliphant and Allen-Vercoe, 2019). Accordingly, the concentration of the SCFAs propionate and isobutyrate were lower in piglets from maternity 1 than in piglets from maternity 2. Since propionate is produced mainly through plant derived polysaccharide degradation (Louis and Flint, 2017), our results suggest that carbohydrate fermentation might be more active in suckling piglets from maternity 2. The different metabolic activity of the microbiota of piglets from the two maternity farms might be explained by their different bacterial composition. Indeed, the relative abundance of two families (Christensenellaceae, Clostridiales vadinBB60) and two genera (*Christensenellaceae R-7 group* and *Ruminococcaceae UCG-002)* were higher in piglets from maternity 2. Members of these bacterial groups are known to be involved in the degradation of plant derived carbohydrates (Biddle *et al.*, 2013; Louis and Flint, 2017; La Reau and Suen, 2018). Overall, our data obtained during the suckling period show that maternal environment influences the microbiota metabolic activity and composition in piglets. More work is needed to determine which factors of the maternal environment drive these effects (e.g. mother diet, creep feed intake, housing, hygiene, animal handling, etc.) (Mulder *et al.*, 2009; Montagne *et al.*, 2010; Megahed *et al.*, 2019).

Despite the effects of weaning on the production of bacterial metabolites were specific to each maternity farm, some of the effects of weaning on the microbiota diversity and composition were observed in piglets from the two maternity farms. There was an increase in OTU richness after weaning, which is in agreement with previous studies in piglets (Frese *et al.*, 2015; Chen *et al.*, 2017; De Rodas *et al.*, 2018). Indeed, the cessation of milk ingestion, the introduction of new plant substrates and the modification of housing environment might contribute to the colonization of the gut by new bacterial species. The relative abundances of Lactobacillaceae and *Lactobacillus* decreased after weaning in piglets from the two maternity farms, as previously observed in numerous studies in piglets (Mach *et al.*, 2015; Slifierz *et al.*, 2015; Bian *et al.*, 2016; Chen *et al.*, 2017; De Rodas *et al.*, 2018; Guevarra *et al.*, 2018). Members of the *Lactobacillus* genus are specialized in the degradation of lactose (W. Wang *et al.*, 2019), the main sugar provided by maternal milk (4.8% in pigs *vs* 6.8% in humans) (Zhang *et al.*, 2018). In contrast, the relative abundances of Ruminococcaceae and *Ruminococcus 2* increased after weaning in piglets from the two maternity farms, as previously observed in other studies (Frese *et al.*, 2015; Bian *et al.*, 2016; Chen *et al.*, 2017). The Ruminococcaceae family is well known to play an important role in the degradation of complex plant polysaccharides (La Reau and Suen, 2018). Thus, the increased abundance of Ruminococcaceae after weaning highlights the adaptation of the microbiota to the supply of plant-derived substrates concomitant with suckling cessation.

The taxonomic changes observed after weaning in piglets from both maternity farms were associated with a major shift in the predicted functionality of the microbiota. Indeed, the relative abundance of 62% of the predicted pathways was altered after weaning. Among them, the relative abundance of the “Fucose degradation” pathway decreased in piglets from the two maternity farms. Fucose is a monosaccharide present in milk oligosaccharides (fucosylated oligosaccharides represent 9.1% of total milk oligosaccharides in sow) (Salcedo *et al.*, 2016). Thus, the predicted decrease in the relative abundance of the “Fucose degradation” pathway might be driven by suckling cessation at weaning. The OTU contributing the most to this predicted pathway was classified in Enterobacteriaceae (dominant family during the suckling period). This result is in agreement with a study using shotgun metagenomics in suckling piglets showing that the majority of genes related to fucose utilization were assigned to Enterobacteriaceae (Salcedo *et al.*, 2016). Another interesting result was the increased relative abundance of the predicted pathway “Starch degradation V” after weaning in piglets from both maternity farms. Starch is an important source of carbohydrates for the microbiota of weaned piglets (W. Wang *et al.*, 2019). Our functional predictions are in agreement with a shotgun metagenomic study showing that bacterial genes coding for starch degrading-enzymes were more abundant after weaning in piglets (Frese *et al.*, 2015). We also found that after weaning the OTU contributing the most to the “Starch degradation V” predicted pathway was assigned to *Ruminoccocaceae*. Accordingly, a member of this family, *Ruminococcus bromii*, was demonstrated to be a key stone species for starch degradation in the human microbiota (Ze *et al.*, 2012). Overall, our results suggest that suckling cessation and the shift to a plant based diet drive the maturation of the microbiota composition and predicted functionality observed after weaning in piglets.

In contrast to the shared effects of weaning on the microbiota composition and predicted functionality observed in piglets from the two maternity farms, the alteration of bacterial metabolites production was specific to piglets from each maternity farm. In piglets from maternity 1, there was a reduction of the relative concentrations of biogenic amines (cadaverine, tyramine and 5-aminovalerate) after weaning. These metabolites are produced by the gut microbiota from amino acids degradation (lysine, tyrosine and proline, respectively) (Barker *et al.*, 1987; Portune *et al.*, 2016). The relevance for gut health of these bacterial biogenic amines is difficult to predict since both protective and toxic effects have been described, mostly depending on the concentration (Louis *et al.*, 2014; Blander *et al.*, 2017; Oliphant and Allen-Vercoe, 2019). The reduction of their concentration after weaning might be linked to the decreased relative abundance of Enterobacteriaceae, members of this family being known to produce biogenic amines (Oliphant and Allen-Vercoe, 2019; Tofalo *et al.*, 2019). The decrease after weaning of the relative abundance of Proteobacteria, Enterobacteriaceae and *Escherichia-Shigella* (11.4% during the suckling period vs 4.6% after weaning) observed in piglets from maternity 1 is in agreement with several previous studies in piglets (Frese *et al.*, 2015; Bian *et al.*, 2016; De Rodas *et al.*, 2018). The high abundance of Enterobacteriaceae in early life can be linked to their adaptation to use substrates available in the gut of suckling mammals. Indeed, some species including members of the *Bacteroides* degrade milk oligosaccharides externally which releases free sugars in the lumen that can promote the growth of Enterobacteriaceae through cross-feeding reactions (Charbonneau *et al.*, 2016). Thus, the reduction after weaning of the relative abundance of Enterobacteriaceae in piglets from maternity 1 might be related to the cessation of suckling. In these piglets, there was also an increase after weaning in the relative concentrations of acetate and propionate. This increase in SCFA concentration after weaning was reported previously in piglets (van Beers-Schreurs *et al.*, 1998) and might have protective effects for gut health due to their capacity to reinforce the mucosal barrier and to support immune functions (Koh *et al.*, 2016). The upregulation of acetate and propionate production by the microbiota at weaning reflects the metabolic adaptation of the gut bacteria to solid feed derived substrates since these SCFAs are produced mainly through plant carbohydrate fermentation (Louis and Flint, 2017). Besides this substrate effects, the increase SCFA production in piglets from maternity 1 after weaning might also be linked to the increase in the relative abundance of the Christensenellaceae, *Christensenellaceae R-7 group* and *Ruminococcaceae UCG-002*, also observed in previous studies (Frese *et al.*, 2015; Bian *et al.*, 2016; Chen *et al.*, 2017). The ability of these bacterial groups to break down complex plant polysaccharides suggests that the shift from maternal milk to a plant based diet favored their growth (La Reau and Suen, 2018). In summary, the changes in microbiota metabolic activity and composition observed at weaning only in piglets from maternity 1 were probably driven by an abrupt dietary shift from milk-derived substrates (e.g. oligosaccharides) to plant-derived substrates.

Strikingly, most of the modification observed after weaning in piglets from maternity 1 were not found in piglets from maternity 2. This might be linked to the more “mature state” of the microbiota composition and metabolic activity observed already during the suckling period in these piglets, as discussed above (e.g. higher relative abundance of *Ruminococcaceae UCG-002* and *Christensenellaceae R-7 group* and higher relative concentration of SCFA). Interestingly, some differences observed during the suckling period (e.g. higher relative abundance of *Ruminococcaceae UCG-002)* were still found after weaning, showing a persistent effect of the maternal environment. Thus, our study highlights the importance of early life environment on the microbiota composition and functionality during the suckling period but also after weaning.

In conclusion, our study shows that the functional maturation of the microbiota at weaning is influenced by early life environment in piglets. Since this alteration of the microbiota functionality at the suckling-to-solid food transition might play a key role in intestinal and immune maturation (Hooper, 2004; Al Nabhani and Eberl, 2020), a perspective of our work would be to determine what are the effects on the gut barrier function of the metabolites produced by piglet microbiota before or after weaning. In this context, the recent development of pig intestinal organoids represent a great opportunity to decipher the action of gut microbiota derived metabolites on piglet epithelial cells (van der Hee *et al.*, 2018). Subsequently, innovative strategies based on the control of early life environment might be developed to orient the metabolic activity of the microbiota towards the production of protective metabolites promoting intestinal development and long-term health.

## EXPERIMENTAL PROCEDURES

### Animals

The experiments were performed in two separate maternity farms (maternity 1 and maternity 2) and one post-weaning farm, all located in Morbihan, France. Piglets (Piétrain x Large white x Landrace) were housed in maternity 1 (n=14 piglets/10 sows) or maternity 2 (n=14 piglets/9 sows) from birth to day 21 (weaning day) (figure 1). Chemical composition of gestation and lactation diets of sows from the two maternity farms are shown in table S1. Suckling piglets had access to creep feed from day 1 in maternity 1 and from day 7 in maternity 2 (chemical composition of piglets diets shown in table S1). At weaning (day 21), piglets from the two maternity farms were moved to a single post-weaning farm and mixed in the same pens in one room. After weaning, all piglets had access to the same diet (table S1). None of the piglets received antibiotic treatment. Fecal samples were collected during the suckling period (day 13) and two days after weaning (day 23) from the same piglets and stored at - 80°C until analysis.

### 16S rRNA gene sequencing and sequences analysis

Fecal DNA was extracted using Quick-DNA Fecal/Soil Microbe 96 Kit (ZymoResearch, Irvine, CA) and the 16S rRNA V3-V4 region was amplified by PCR and sequenced by MiSeq Illumina Sequencing as previously described (Verschuren *et al.*, 2018). Sequencing reads were deposited in the National Center for Biotechnology Information Sequence Read Archive (SRA accession: PRJNA591810). 16S rDNA amplicon sequences were analyzed using the FROGS pipeline according to standard operating procedures (Escudié *et al.*, 2018). Amplicons were filtered according to their size (400-500 nucleotides) and clustered into OTUs using Swarm (aggregation distance: d=1 + d=3). After chimera removal, OTUs were kept when representing more than 0.005% of the total number of sequences (Bokulich *et al.*, 2013). OTUs affiliation was performed using the reference database silva132 16S with a minimum pintail quality of 80 (Quast *et al.*, 2013). The mean number of reads per sample was 15 639 (min: 9 694 - max: 30 724). The functional potential of the microbiota was predicted by using PICRUSt2 (Douglas *et al.*, 2019) according to the guidelines with the unrarefied OTU abundance table as input. Relative predicted abundance of MetaCyc pathways were calculated by dividing the abundance of each pathway by the sum of all pathways abundances per sample. Relative contribution of each OTU to predicted pathways was calculated by dividing the contribution of each OTU by the sum of all contributions per sample.

### NMR metabolomics

Feces (100 mg) were homogenized in 500 µL phosphate buffer (prepared in D2O, pH7, TSP 1 mM) in 2mL FastPrep tubes (Lysing D matrix) by using a FastPrep Instrument (MP biomedicals, Irvine, CA). After centrifugation (12 000 g, 4°C, 10 min), supernatants were collected. The extraction step was repeated on the pellet. Supernatants were pooled and centrifuged twice (18 000 g, 30 min, 4°C). The resulting supernatant (600 µL) was transferred to a 5 mm NMR tube. All NMR spectra were obtained with an Avance III HD NMR spectrometer operating at 600.13 MHz for ^1^H resonance frequency using a 5 mm inverse detection CryoProbe (Bruker Biospin, Rheinstetten, Germany) in the MetaboHUB-MetaToul-AXIOM metabolomics platform (Toulouse, France). ^1^H NMR spectra were acquired at 300 K using the Carr-Purcell-Meiboom-Gill spin-echo pulse sequence with presaturation. Pre-processing of the spectra (group delay correction, solvent suppression, apodization with a line broadening of 0.3 Hz, Fourier transform, zero order phase correction, shift referencing on TSP, baseline correction, setting of negative values to zero) was performed in the galaxy tool Workflow4Metabolomics following guidelines (Giacomoni *et al.*, 2015). After water region (4.5 – 5.1 ppm) exclusion, spectra (0.5 – 9 ppm) were bucketed (0.01 ppm bucket width) and normalized by total area in Workflow4Metabolomics. Representative samples were characterized by 2D NMR experiments (^1^H-^1^H COSY and ^13^C-^1^H HSQC). For metabolite identification, 1D and 2D NMR spectra of pure compounds prepared in the same buffer and acquired with the same spectrometer were overlayed with samples spectra. Annotated representative spectra are presented in figure 2. For each identified metabolite, buckets non-overlapping with other metabolites were selected for the quantification (table 1).

### Statistical analysis

All statistical analysis were performed using the R software (version 3.5.1). PLS-DA was performed with the mixOmics package (Rohart *et al.*, 2017). Metabolites relative concentrations were used as variable matrix (X). Experimental groups (defined by day and maternity) were used as predictors (Y). The microbiota composition analysis was performed using the phyloseq package (McMurdie and Holmes, 2013). For α and β diversity analyses, the samples were rarefied to even sequencing depth (9 694 reads per sample). Richness (observed OTUs), Shannon and Inverse Simpson α-diversity indices were calculated. The β-diversity was analyzed using the Bray-Curtis distance and plotted by non-Metric Dimensional Scaling (nMDS). Bacterial taxa differential abundance analysis was performed with unrarefied data. OTUs representing less than 0.05% of the total number of sequences were filtered out. OTUs were agglomerated at phylum, family or genus level and relative abundances were calculated at each taxonomic level. Heatmaps and clustering analysis (average method) were performed with the made4 package. All univariate analyses were performed with a linear mixed model (lme4 package) with the fixed effects of age (day 13 or day 23), maternity (maternity 1 or 2) and their interaction, while piglet and sow were used as random effect. The relative abundances of bacterial groups, the relative abundances of predicted pathways and the relative contributions of OTUs to predicted pathways were log transformed. Analysis of variance was performed with the car package and P values were adjusted for multiple testing with the false discovery rate (FDR) procedure. Means of each group were compared pairwise with the emmeans package (Tukey correction). P<0.05 were considered significant.

## ACKNOWLEDGMENTS

The authors are grateful to the genotoul bioinformatics platform Toulouse Occitanie (Bioinfo Genotoul, doi: 10.15454/1.5572369328961167E12) and Sigenae group for providing computing and storage resources to Galaxy instance (http://sigenae-workbench.toulouse.inra.fr).

## AUTHOR CONTRIBUTIONS

AS and SC conceived the experiments. MB, IA, CB, OB performed the experiments. MB, LC, AB, CC, OZ, GP, AS, SC analyzed the data. MB, AS and SC wrote the manuscript.

## SUPPLEMENTARY MATERIALS

**Table S1: Chemical composition of sows and piglets diets.**

**Table S2: Univariate statistical analysis of metabolites relative concentrations in piglet feces.** Metabolome was analyzed by NMR-based metabolomics in feces collected from piglets from two maternity farms (M1 and M2) during the suckling period (D13) and after weaning (D23). Columns B to D: For each metabolite (row), a linear mixed model was used with age and maternity as fixed effects and sows and piglets as random effects. P-values were adjusted with the false discovery rate (FDR) method. P-values < 0.05 are indicated in bold. Columns E to G: Non adjusted P-values. Columns H to K: Means were compared pairwise and p-values were adjusted with the Tukey method. The means of groups associated with different letters are significantly different (P<0.05). Columns L to O: The mean relative concentration of each metabolite is presented for each group.

**Table S3: Univariate statistical analysis of microbiota α-diversity in piglet feces.** The microbiota composition was analyzed by 16S rRNA gene in feces collected from piglets from two maternity farms (M1 and M2) during the suckling period (D13) and after weaning (D23). Columns B to D: For each α-diversity index (row), a linear mixed model was used with age and maternity as fixed effects and sows and piglets as random effects. P-values < 0.05 are indicated in bold. Columns E to H: Means were compared pairwise and p-values were adjusted with the Tukey method. The means of groups associated with different letters are significantly different (P<0.05). Columns I to L: The mean of each α-diversity index is presented for each group.

**Table S4: Univariate statistical analysis of bacterial phyla relative abundance in piglet feces.** The microbiota composition was analyzed by 16S rRNA gene in feces collected from piglets from two maternity farms (M1 and M2) during the suckling period (D13) and after weaning (D23). Columns B to D: For each phylum (row), a linear mixed model was used with age and maternity as fixed effects and sows and piglets as random effects. P-values were adjusted with the false discovery rate (FDR) method. P-values < 0.05 are indicated in bold. Columns E to G: Non adjusted P-values. Columns H to K: Means were compared pairwise and p-values were adjusted with the Tukey method. The means of groups associated with different letters are significantly different (P<0.05). Columns L to O: The mean relative abundance of each phylum is presented for each group.

**Table S5: Univariate statistical analysis of bacterial families relative abundance in piglet feces.** The microbiota composition was analyzed by 16S rRNA gene in feces collected from piglets from two maternity farms (M1 and M2) during the suckling period (D13) and after weaning (D23). Columns B to D: For each family (row), a linear mixed model was used with age and maternity as fixed effects and sows and piglets as random effects. P-values were adjusted with the false discovery rate (FDR) method. P-values < 0.05 are indicated in bold. Columns E to G: Non adjusted P-values. Columns H to K: Means were compared pairwise and p-values were adjusted with the Tukey method. The means of groups associated with different letters are significantly different (P<0.05). Columns L to O: The mean relative abundance of each family is presented for each group.

**Table S6: Univariate statistical analysis of bacterial genera relative abundance in piglet feces.** The microbiota composition was analyzed by 16S rRNA gene in feces collected from piglets from two maternity farms (M1 and M2) during the suckling period (D13) and after weaning (D23). Columns B to D: For each genus (row), a linear mixed model was used with age and maternity as fixed effects and sows and piglets as random effects. P-values were adjusted with the false discovery rate (FDR) method. P-values < 0.05 are indicated in bold. Columns E to G: Non adjusted P-values. Columns H to K: Means were compared pairwise and p-values were adjusted with the Tukey method. The means of groups associated with different letters are significantly different (P<0.05). Columns L to O: The mean relative abundance of each genus is presented for each group.

**Table S7: Univariate statistical analysis of OTUs relative abundance in piglet feces.** The microbiota composition was analyzed by 16S rRNA gene in feces collected from piglets from two maternity farms (M1 and M2) during the suckling period (D13) and after weaning (D23). Columns E to G: For each OTU (row), a linear mixed model was used with age and maternity as fixed effects and sows and piglets as random effects. P-values were adjusted with the false discovery rate (FDR) method. P-values < 0.05 are indicated in bold. Columns H to J: Non adjusted P-values. Columns K to N: Means were compared pairwise and p-values were adjusted with the Tukey method. The means of groups associated with different letters are significantly different (P<0.05). Columns O to R: The mean relative abundance of each OTU is presented for each group.

**Table S8: Univariate statistical analysis of predicted bacterial pathways relative abundance in piglet feces.** The potential functionality of the gut microbiota was inferred from 16S rRNA amplicon sequences in feces collected from piglets from two maternity farms (M1 and M2) during the suckling period (D13) and after weaning (D23). Columns B to D: For each predicted pathway (row), a linear mixed model was used with age and maternity as fixed effects and sows and piglets as random effects. P-values were adjusted with the false discovery rate (FDR) method. P-values < 0.05 are indicated in bold. Columns E to G: Non adjusted P-values. Columns H to K: Means were compared pairwise and p-values were adjusted with the Tukey method. The means of groups associated with different letters are significantly different (P<0.05). Columns L to O: The mean relative abundance of each predicted pathway is presented for each group.

**Table S9: Univariate statistical analysis of the relative contribution of each OTU to the pathway “Fucose degradation”**. The relative contribution of each OTU to the predicted pathway was inferred from 16S rRNA amplicon sequences in feces collected from piglets from two maternity farms (M1 and M2) during the suckling period (D13) and after weaning (D23). Columns B to D: For each OTU (row), a linear mixed model was used with age and maternity as fixed effects and sows and piglets as random effects. P-values were adjusted with the false discovery rate (FDR) method. P-values < 0.05 are indicated in bold. Columns E to G: Non adjusted P-values. Columns H to K: Means were compared pairwise and p-values were adjusted with the Tukey method. The means of groups associated with different letters are significantly different (P<0.05). Columns L to O: The mean relative contribution of each OTU is presented for each group.

**Table S10: Univariate statistical analysis of the relative contribution of each OTU to the pathway “Starch degradation V”**. The relative contribution of each OTU to the predicted pathway was inferred from 16S rRNA amplicon sequences in feces collected from piglets from two maternity farms (M1 and M2) during the suckling period (D13) and after weaning (D23). Columns B to D: For each OTU (row), a linear mixed model was used with age and maternity as fixed effects and sows and piglets as random effects. P-values were adjusted with the false discovery rate (FDR) method. P-values < 0.05 are indicated in bold. Columns E to G: Non adjusted P-values. Columns H to K: Means were compared pairwise and p-values were adjusted with the Tukey method. The means of groups associated with different letters are significantly different (P<0.05). Columns L to O: The mean relative contribution of each OTU is presented for each group.

